# Reduced effectiveness of purifying selection on new mutations in a parthenogenic terrestrial isopod (*Trichoniscus pusillus*)

**DOI:** 10.1101/2023.01.03.522635

**Authors:** Emily Yarbrough, Christopher H. Chandler

## Abstract

The “paradox of sex” refers to the question of why sexual reproduction is maintained in the wild, despite how costly it is compared to asexual reproduction. Because of these costs, one would suspect nature would select for asexual reproduction, yet sex seems to be continually selected for. Multiple hypotheses have been proposed to explain this incongruence, including the niche differentiation hypothesis, the Red Queen hypothesis, and accumulation of harmful mutations in asexual species due to inefficient purifying selection. This study focuses on the accumulation of mutations in the closely related terrestrial isopod species *Trichoniscus pusillus*, which has sexual diploid and parthenogenic triploid forms, and *Hyloniscus riparius*, an obligately sexual relative. We surveyed sex ratios of both species in an upstate New York population and obtained RNA-seq data from wild-caught individuals of both species to examine within- and between-species patterns of molecular evolution in protein-coding genes. The sex ratio and RNA-seq data together provide strong evidence that this *T. pusillus* population is entirely asexual and triploid, while the *H. riparius* population is sexual and diploid. Although all wild-caught *T. pusillus* individuals used for sequencing shared identical genotypes at nearly all SNPs, supporting a clonal origin for this population, heterozygosity and SNP density were much higher in *T. pusillus* than in the sexually reproducing *H. riparius*, suggesting this parthenogenic lineage may have arisen via mating between two divergent diploid lineages. Between-species sequence comparisons showed no evidence of ineffective purifying selection in the asexual *T. pusillus* lineage, as measured by the ratio of nonsynonymous to synonymous substitutions (dN/dS ratios). Likewise, there was no difference between *T. pusillus* and *H. riparius* in the ratios of nonsynonyous to synonymous SNPs overall (pN/pS). However, pN/pS ratios in *T. pusillus* were significantly higher when considering only SNPs that may have arisen via recent mutation after the transition to parthenogenesis. Thus, these recent SNPs support the hypothesis that purifying selection is less effective against new mutations in asexual lineages, but that this only becomes apparent over long time scales. This system provides a useful model for future studies on the evolutionary tradeoffs between sexual and asexual reproduction in nature.

**Significance:** Asexual organisms are expected to have several key evolutionary advantages, most multicellular organisms reproduce sexually. One hypothesis to explain why sex is so common despite its costs proposes that recombination allows selection to remove harmful mutations from sexual populations more efficiently. We tested this hypothesis by looking at sequence variants in expressed genes in a novel study system, an asexual terrestrial isopod crustacean and a close sexual relative. Our results support the hypothesis that selection is less effective at filtering out harmful mutations in asexual organisms, but that longer time scales may be needed for this cost of asexual reproduction to become substantial.

## Introduction

The “paradox of sex”, the question of why sex is maintained in the wild, is a topic of great debate among evolutionary biologists (Neiman et al. 2018). Theoretically speaking, nature should select for asexual reproduction, as sexual reproduction requires far more resources. For example, a sexually reproducing organism must allocate resources toward locating a mate, courtship, etc. (Otto 2009). There are also overall costs associated with the production of males and recombination (Lehtonen et al. 2012; Gibson et al. 2017). However, sexual reproduction is incredibly prevalent in eukaryotic organisms, suggesting it is maintained by selection (Otto 2009; Speijer et al. 2015). This contradiction has been an ongoing topic of debate for years, and multiple hypotheses have been proposed to explain the prevalence of sex. Some of the more famous examples include the niche differentiation hypothesis, the Red Queen Hypothesis, and the accumulation of harmful mutations within the genomes of the asexual species (Neiman et al. 2018). The niche differentiation hypothesis proposes that sexually reproducing species occupy broader ecological niches or have less niche overlap with asexuals, reducing competition between sexuals and asexuals and counteracting the costs of sex (Neiman et al. 2018). The Red Queen hypothesis suggests that host-parasite co-evolution selects for sexual recombination (Agrawal 2009). In this situation, it would be more beneficial to be a sexually reproducing organism as negative-frequency dependent selection would favor rare host genotypes (Lively 1987; Agrawal 2009; Peters & Lively 1999).

Sexual organisms also have the benefit of segregation and recombination to facilitate efficient purifying selection, purging deleterious mutations from the population (Felsenstein 1974; MacPherson et al. 2021; Kondrashov 1994; Hartfield & Keightley 2012; Hollister et al. 2015). Recombination is especially effective at purging such mutations as it not only prevents disadvantageous mutations from becoming established in a population, but it also makes it more likely for favorable mutations to become fixed within a population (Felsenstein 1974). Asexual organisms do not have this benefit, and selection is therefore less effective at removing any deleterious mutations from their genomes (Keightley & Otto 2006). In this instance, one could expect asexual species to suffer from mutational meltdown because, as each generation reproduces, there would be continued accumulation of harmful mutations within the population, potentially leading to population decline and extinction (Muller 1964; Lynch et al. 1993).

The idea that an asexually reproducing species will accumulate a larger number of deleterious mutations than their sexual counterparts is generally accepted by biologists (Brandt et al. 2019). However, empirical studies on the topic have provided mixed results. One such study estimated the nonsynonymous to synonymous divergence ratio (dN/dS), a proxy for the effectiveness of purifying selection, in eight sexual and asexual hexapod lineages and did not find evidence suggesting that the asexual species had accumulated more nonsynonymous mutations in their genomes (Brandt et al. 2019). Likewise, transcriptome-based studies in mites (Brandt et al. 2017) and ribbon worms (Ament-Velásquez et al. 2016) failed to support these predictions. On the other hand, in a recent study of asexual *Timema* stick insects, there was evidence for an elevated dN/dS ratio within the asexual populations (Bast et al. 2018). Several other studies mirror this support (Hollister et al. 2015; Johnson & Howard 2007; Neiman et al. 2010; Henry et al. 2012; Maldonado et al. 2022; Lovell et al. 2017), but many of these are based on mitochondrial sequences or a small handful of nuclear genes, rather than genome-wide datasets. Therefore, while the hypothesis of relaxed purifying selection in asexual species is widely accepted, in practice it has met with conflicting results.

To provide a novel test of the hypothesis that inefficient purifying selection in asexuals contributes to the prevalence of sexual reproduction in nature, we examined the terrestrial isopods species *Trichoniscus pusillus* and *Hyloniscus riparius*. Isopods are a relatively novel system for research into the tradeoffs between asexual and sexual reproduction and therefore provide an exciting opportunity for study. *Trichoniscus pusillus* was chosen specifically because it is known to be present in both diploid sexual and triploid asexual populations in Europe (Christensen 1983). *Hyloniscus riparius* was chosen for this study as a potential obligately sexual outgroup for comparison in the same taxonomic family (Schultz 1965; Lins et al. 2017).

This study had two overall goals: 1) determine the sex ratios of our local *T. pusillus* and *H. riparius* populations in Oswego, New York, to establish whether they are sexual or asexual, and 2) investigate patterns of molecular evolution to test the hypothesis that harmful mutations may accumulate in asexual species due to inefficient purifying selection. These goals were achieved by surveying local populations and performing RNA-seq to assemble transcriptomes and identify SNPs for each species to examine patterns of nonsynonymous and synonymous divergence and polymorphism.

## Results

### Field sex ratios

We surveyed *T. pusillus* and *H. riparius* at Rice Creek Field Station in Oswego, NY during summer 2021. All the captured *T. pusillus* individuals were females, whereas *H. riparius* had a mixed sex ratio (Table 1) which was not statistically distinguishable from 1:1 (p = 0.549), albeit with a small sample size. The lack of males in *T. pusillus* suggests this population either reproduces entirely asexually or harbors a sex ratio distorter such as *Wolbachia* (Cordaux et al. 2011). The even sex ratio in *H. riparius* is consistent with a diploid sexual population.

**Table 1.** Observed sex ratio data for *T. pusillus* and *H. riparius* specimens caught in Oswego, NY during summer 2021.

|  | Female | Male | Total |
| --- | --- | --- | --- |
| *T. pusillus* | 170 | 0 | 170 |
| *H. riparius* | 7 | 4 | 11 |

### Transcriptome assembly, SNPs, and heterozygosity

We performed obtained RNA-seq data from four randomly chosen wild-caught adult female *T. pusillus* individuals, one pooled set of offspring from one of these adult females, and four randomly chosen adult *H. riparius* individuals (two males and two females). The sequencing reads obtained for this study have been deposited at the NCBI Sequence Read Archive (BioProject Accession PRJNA916870; SRA Accession Numbers SRR22938942-SRR22938950). As an additional outgroup, we also used RNA-seq reads from two wild-caught adults of *Trachelipus rathkei*, also from Oswego, NY, previously obtained from another another study (Becking et al. 2017) (SRA Accession Numbers SRR5198726 and SRR5198727).

We obtained transcriptome assemblies for each species, with high BUCO completeness scores (< 6% BUSCOs missing for all assemblies; Table 2). We identified thousands of putative SNPs in each species (Tables 2 - 3). Genetic similarity was high in *T. pusillus*, with wild-caught adults, as well as the mother/pooled-offspring pair, sharing identical genotypes at nearly all putative biallelic SNPs (Table 3). Because of the complications inherent to inferring SNP genotypes from transcriptome data, some of the putative SNPs may be false positives due to alternative splicing, recently diverged paralogs, or mapping errors; consistent with this, restricting the analysis to SNPs in transcripts identified as complete, single-copy BUSCO markers resulted in slightly higher genetic similarity among samples in *T. pusillus* (Table 3). In *H. riparius* and *T. rathkei*, on the other hand, a much smaller fraction of SNPs showed identical genotypes among all sequenced samples, whether all SNPs or only SNPs in single-copy BUSCOs were counted (Table 3). Because the number of sequenced samples differed across species, we also counted the fraction of SNPs showing identical genotypes for a random pair of wild-caught adults in each species and observed similar patterns supporting clonality in *T. pusillus* but not *H. riparius* or *T. rathkei* (Table 3).

**Table 2.** Transcriptome assembly statistics for the three species examined in this study. “Main transcripts” refers to the primary transcripts predicted by EvidentialGenes; “alternate transcripts” refers to predicted alternative splice variants of the main transcripts; and “all transcripts” refers to the merged set of main and alternate transcripts. Only SNPs in the main transcripts were counted in the last row (i.e., alternate transcripts were excluded), to avoid potential pseudo-replication caused by SNPs found in multiple splice variants of the same transcript.

|  | *T. pusillus* | *H. riparius* | *T. rathkei* |
| --- | --- | --- | --- |
| Main transcripts | 34,920<br>(35.2 Mb) | 25,408<br>(34.7 Mb) | 33,881<br>(38.8 Mb) |
| Alternate transcripts | 58,041<br>(64.9 Mb) | 36,684<br>(63.7 Mb) | 82,444<br>(114.3 Mb) |
| Total transcripts | 92,961<br>(100.1 Mb) | 62,092<br>(98.4 Mb) | 116,325<br>(153.1 Mb) |
| Complete single-copy BUSCOs (all transcripts) | 248 (24.5%) | 243 (24.0%) | 148 (14.6%) |
| Complete duplicated BUSCOs (all transcripts) | 705 (69.6%) | 757 (74.7%) | 844 (83.3%) |
| Fragmented BUSCOs (all transcripts) | 32 (3.2%) | 8 (0.8%) | 10 (1.0%) |
| Missing BUSCOs (all transcripts) | 28 (2.7%) | 5 (0.5%) | 11 (1.1%) |
| Complete single-copy BUSCOs (main transcripts) | 879 (86.8%) | 929 (91.7%) | 935 (92.3%) |
| Complete duplicated BUSCOs (main transcripts) | 37 (3.7%) | 37 (3.7%) | 37 (3.7%) |
| Fragmented BUSCOs (main transcripts) | 40 (3.9%) | 13 (1.3%) | 17 (1.7%) |
| Missing BUSCOs (main transcripts) | 57 (5.6%) | 34 (3.3%) | 24 (2.3%) |
| Total SNPs / biallelic SNPs with genotype data in all samples (number of samples) | 134,485 /<br>5,401 (5) | 53,613 /<br>12,676 (4) | 32,542 /<br>17,684 (2) |

**Table 3.** Genetic similarity among individuals as assessed by identical genotypes at SNPs in transcriptome sequencing data. In this analysis, SNPs showing evidence of more than two alleles were excluded.

|  | *T. pusillus* | *H. riparius* | *T. rathkei* |
| --- | --- | --- | --- |
| All biallelic SNPs | Total SNPs: 5,401<br>All adults identical: 4,961 (91.9%)<br>Mother-offspring identical: 4,905 (90.8%)<br>Two adults identical: 5,201 (96.5%) | Total SNPs: 12,676<br>All adults identical: 1,208 (9.5%)<br>Two adults identical: 7,081 (55.9%) | Total SNPs: 17,684<br>Both adults identical: 4,983 (28.2%) |
| Biallelic SNPs in single-copy BUSCO transcripts | Total SNPs: 465<br>All adults identical: 452 (97.2%)<br>Mother-offspring identical: 439 (94.4%)<br>Two adults identical: 458 (98.5%) | Total SNPs: 1,054<br>All adults identical: 26 (2.5%)<br>Two adults identical: 625 (59.3%) | Total SNPs: 1,078<br>Both adults identical: 308 (28.6%) |

To assess ploidy levels, the proportion of reads containing the reference allele vs. the alternate allele in heterozygous individuals was estimated for all SNPs in each species (Figure 1). For *T. pusillus* individuals, there are two noticeable peaks in the frequency of reference read counts at ∼0.33 and ∼0.67; the peak at 0.33 is much higher than the peak at 0.67, perhaps because some assemblers are biased with respect to which allele is selected as the “reference” allele. For *H. riparius* and *T. rathkei*, there is only one noticeable peak in the frequency of alleles near the ∼0.5 mark, although the peak is slightly below the 0.5 mark in both species, perhaps because of biases in SNP detection or alignment. Regardless, these patterns are consistent with triploidy in *T. pusillus* and diploidy in *H. riparius*, further supporting the hypothesis that triploid parthenogenesis is the primary reproductive mode in this population of *T. pusillus*.

**Figure 1.**
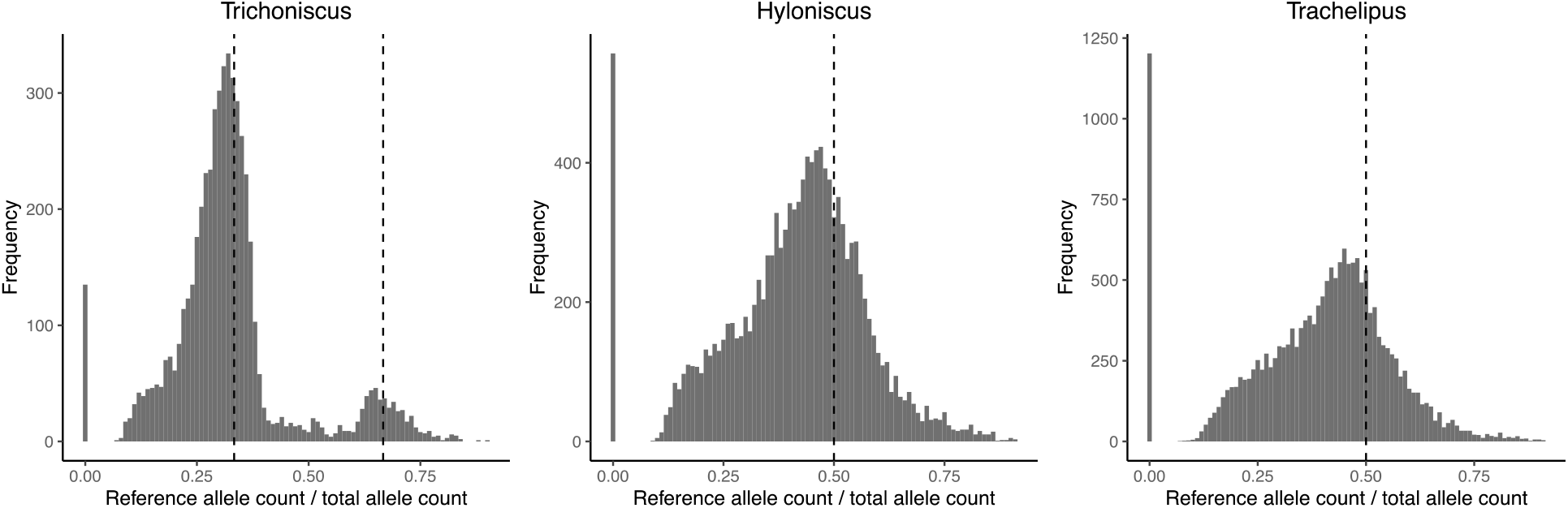
Histogram illustrating the distribution of the proportion of reads containing the reference allele (as opposed to the alternate allele) at each SNP in heterozygous individuals within each species. In *T. pusillus*, there are two peaks at approximately 0.33 and 0.66, consistent with triploid genotypes. In *H. riparius*, there is one peak at approximately 0.5, consistent with diploid genotypes.

To investigate heterozygosity levels among species, the proportion of heterozygous nucleotide sites within each individual was estimated for *T. pusillus, H. riparius*, and the outgroup *Trachelipus rathkei* (Table 4). *T. pusillus* individuals displayed roughly 0.7-1.0% heterozygosity, as compared to the 0.13-0.25% heterozygosity of *H. riparius* and 0.21-0.26% heterozygosity of the *Trachelipus* outgroup. The elevated heterozygosity in triploid *T. pusillus* suggests it may have arisen via hybridization between divergent lineages.

**Table 4.**
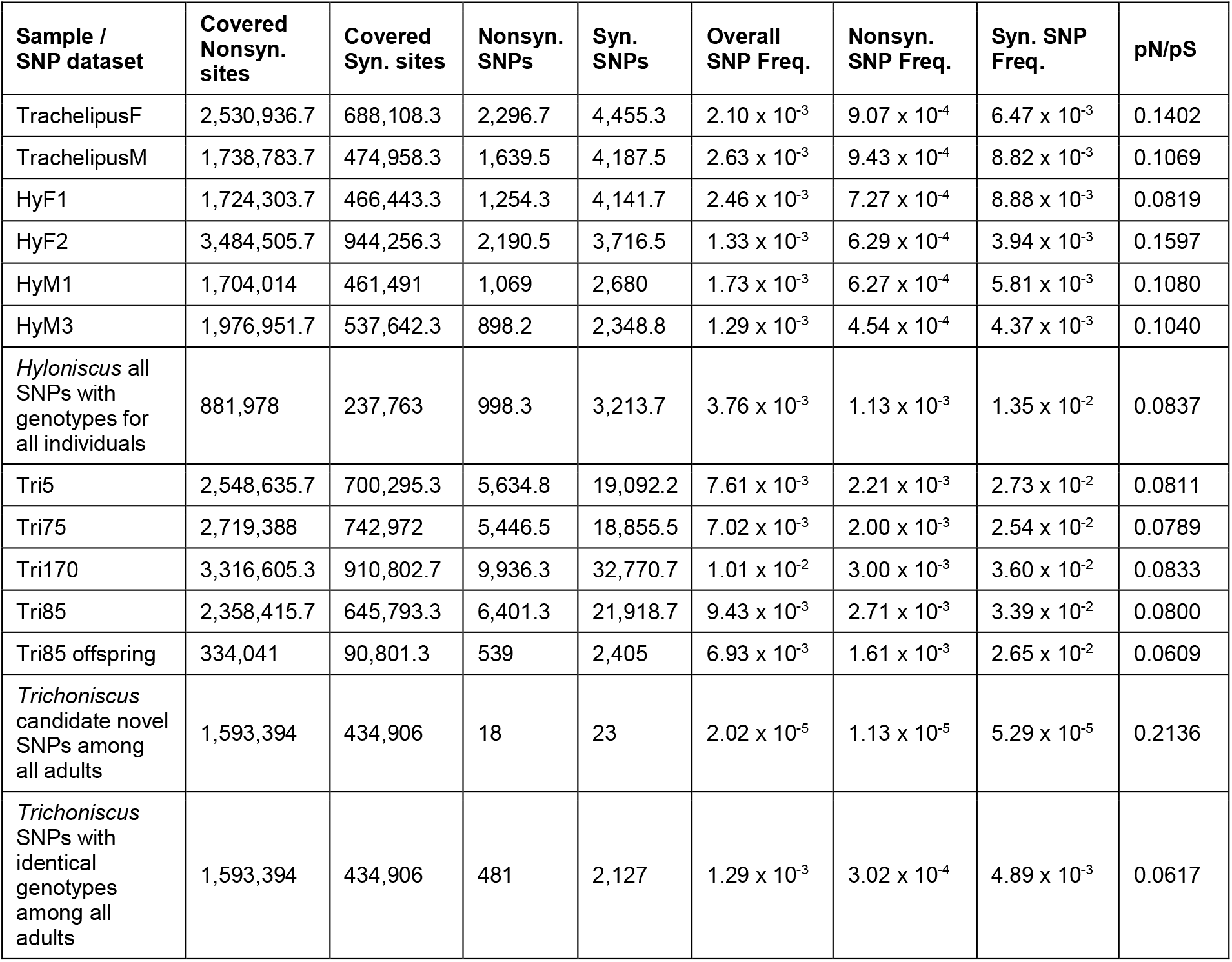
Within-species pN/pS ratios for heterozygous SNPs within each individual, or SNPs across multiple samples. These numbers only include SNPs within predicted full-length coding sequences. Covered sites refer to nucleotide sites with sequencing depth of at least 20 in that individual, or depth of at least 20 in every individual for SNPs across multiple samples.

| Sample / SNP dataset | Covered Nonsyn. sites | Covered Syn. sites | Nonsyn. SNPs | Syn. SNPs | Overall SNP Freq. | Nonsyn. SNP Freq. | Syn. SNP Freq. | pN/pS |
| --- | --- | --- | --- | --- | --- | --- | --- | --- |
| TrachelipusF | 2,530,936.7 | 688,108.3 | 2,296.7 | 4,455.3 | $2.10 \times 10^{-3}$ | $9.07 \times 10^{-4}$ | $6.47 \times 10^{-3}$ | 0.1402 |
| TrachelipusM | 1,738,783.7 | 474,958.3 | 1,639.5 | 4,187.5 | $2.63 \times 10^{-3}$ | $9.43 \times 10^{-4}$ | $8.82 \times 10^{-3}$ | 0.1069 |
| HyF1 | 1,724,303.7 | 466,443.3 | 1,254.3 | 4,141.7 | $2.46 \times 10^{-3}$ | $7.27 \times 10^{-4}$ | $8.88 \times 10^{-3}$ | 0.0819 |
| HyF2 | 3,484,505.7 | 944,256.3 | 2,190.5 | 3,716.5 | $1.33 \times 10^{-3}$ | $6.29 \times 10^{-4}$ | $3.94 \times 10^{-3}$ | 0.1597 |
| HyM1 | 1,704,014 | 461,491 | 1,069 | 2,680 | $1.73 \times 10^{-3}$ | $6.27 \times 10^{-4}$ | $5.81 \times 10^{-3}$ | 0.1080 |
| HyM3 | 1,976,951.7 | 537,642.3 | 898.2 | 2,348.8 | $1.29 \times 10^{-3}$ | $4.54 \times 10^{-4}$ | $4.37 \times 10^{-3}$ | 0.1040 |
| *Hyloniscus* all SNPs with genotypes for all individuals | 881,978 | 237,763 | 998.3 | 3,213.7 | $3.76 \times 10^{-3}$ | $1.13 \times 10^{-3}$ | $1.35 \times 10^{-2}$ | 0.0837 |
| Tri5 | 2,548,635.7 | 700,295.3 | 5,634.8 | 19,092.2 | $7.61 \times 10^{-3}$ | $2.21 \times 10^{-3}$ | $2.73 \times 10^{-2}$ | 0.0811 |
| Tri75 | 2,719,388 | 742,972 | 5,446.5 | 18,855.5 | $7.02 \times 10^{-3}$ | $2.00 \times 10^{-3}$ | $2.54 \times 10^{-2}$ | 0.0789 |
| Tri170 | 3,316,605.3 | 910,802.7 | 9,936.3 | 32,770.7 | $1.01 \times 10^{-2}$ | $3.00 \times 10^{-3}$ | $3.60 \times 10^{-2}$ | 0.0833 |
| Tri85 | 2,358,415.7 | 645,793.3 | 6,401.3 | 21,918.7 | $9.43 \times 10^{-3}$ | $2.71 \times 10^{-3}$ | $3.39 \times 10^{-2}$ | 0.0800 |
| Tri85 offspring | 334,041 | 90,801.3 | 539 | 2,405 | $6.93 \times 10^{-3}$ | $1.61 \times 10^{-3}$ | $2.65 \times 10^{-2}$ | 0.0609 |
| *Trichoniscus* candidate novel SNPs among all adults | 1,593,394 | 434,906 | 18 | 23 | $2.02 \times 10^{-5}$ | $1.13 \times 10^{-5}$ | $5.29 \times 10^{-5}$ | 0.2136 |
| *Trichoniscus* SNPs with identical genotypes among all adults | 1,593,394 | 434,906 | 481 | 2,127 | $1.29 \times 10^{-3}$ | $3.02 \times 10^{-4}$ | $4.89 \times 10^{-3}$ | 0.0617 |

### dN/dS and pN/pS

We identified 11,136 putative one-to-one (single-copy) orthologs across the three species. We estimated dN/dS ratios using pairwise comparisons, between *T. pusillus* and *T. rathkei*, and between *H. riparius* and *T. rathkei* (Figure 2). Across these orthologs, *T. pusillus* had significantly higher estimates of dN (0.161 and 0.152 median values for *T. pusillus* and *H. riparius*, respectively; Wilcoxon paired sample signed rank test p = 1.4 × 10^−69^) and dS (7.22 and 6.60 median values for *T. pusillus* and *H. riparius*, respectively; Wilcoxon paired sample signed rank test p = 4.0 × 10^−11^) but significantly lower estimates of dN/dS (median estimates of 0.019 and 0.021 for *T. pusillus* and *H. riparius*, respectively; Wilcoxon paired sample signed rank test p = 1.2 × 10^−6^). However, the differences appear small in magnitude, and the distributions of dN, dS, and dN/dS overlap almost entirely for the two species (Figure 2). These trends were robust to the choice of codon substitution model (codon frequencies assumed to be equal, CodonFreq=0; codon frequencies estimated from average nucleotide frequencies, CodonFreq=1; codon frequencies estimated from average nucleotide frequencies at the three codon positions, CodonFreq=2; codon frequencies included as free parameters, CodonFreq=3), although the exact estimates of dN, dS, and dN/dS varied slightly as expected (Supplementary Figure 1).

**Figure 2.**
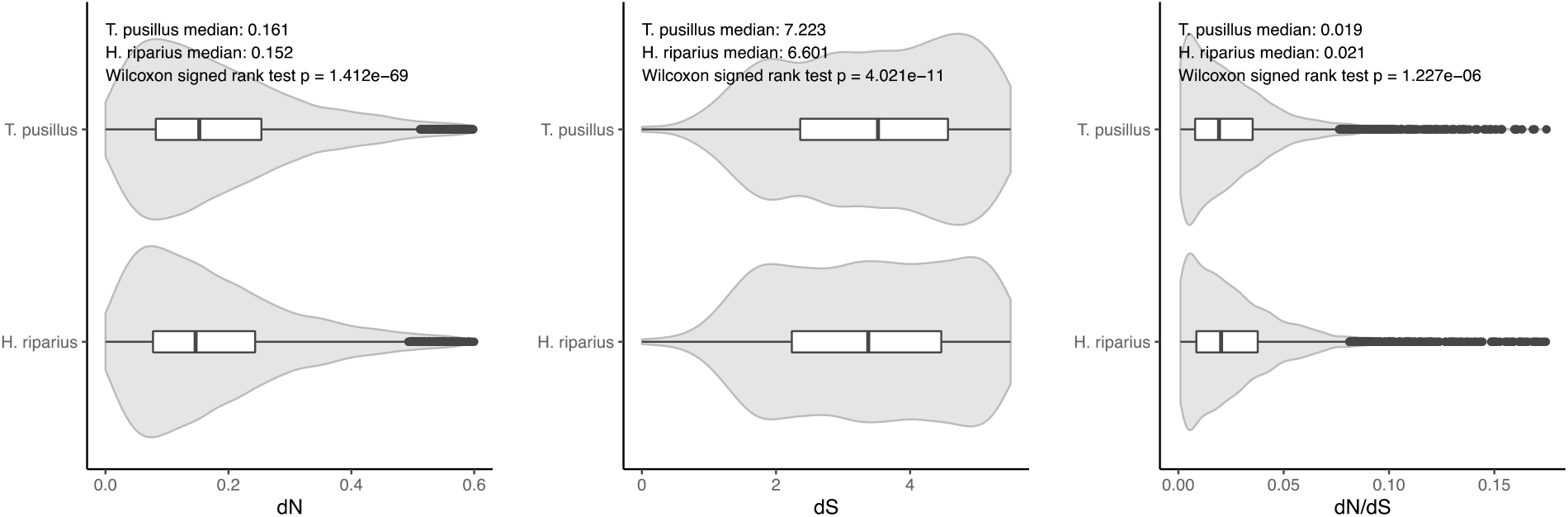
Violin plots of *dN, dS*, and *dN/dS* ratios, resulting from pairwise comparisons between *Trichoniscus* vs. *Trachelipus*, and *Hyloniscus* vs. *Trachelipus*.

At heterozygous sites within individuals in *T. pusillus* and *H. riparius*, there was no significant difference between species in pN/pS, the ratio of non-synonymous to synonymous polymorphisms at heterozygous sites in individual samples (Table 4; p = 0.057, unpaired Wilcoxon signed rank test), although overall rates of heterozygosity were higher in *T. pusillus* as described above; if anything, pN/pS ratios at heterozygous sites within individuals were slightly lower in *T. pusillus* than in *H. riparius*. However, given that the high number of heterozygous sites in *T. pusillus* may be the result of a cross between divergent lineages at or just prior to the transition to parthenogenesis, we sought to examine a subset of SNPs that may represent recent mutations, after the origin of parthenogenesis. While the *T. pusillus* individuals in our population shared identical genotypes at nearly all SNPs, suggesting a recent clonal origin for this lineage, we might expect genotypes to differ among individuals at SNPs that arose recently within this population. Therefore, we searched for SNPs where wild-caught adult *T. pusillus* individuals (excluding the pooled offspring) had different genotypes, such that some individuals were homozygous for one, putatively ancestral, allele, and other individuals carried one copy of a putative derived “mutant” allele. In other words, we filtered for SNPs at least one individual had the genotype 0/0/0 and the others had the genotype 0/0/1, with 0 and 1 representing the putative ancestral and derived alleles, respectively. We then looked at the ratio of nonsynonymous to synonymous polymorphisms in this set of candidate recent SNPs. The pN/pS ratio at these SNPs, putatively arising from recent mutations, was significantly higher than the pN/pS ratio at SNPs in *T. pusillus* where all individuals had identical genotypes (χ^2^ = 17.1; df = 1; p = 0.000035), and higher than overall pN/pS ratios in *H. riparius* (χ^2^ = 9.1; df = 1; p = 0.0025).

## Discussion

### Apomictic parthenogenesis and triploidy in T. pusillus

This population of *T. pusillus* in Oswego, NY, is likely asexual, with clonal, apomictic reproduction, based on multiple lines of evidence. First, all specimens gathered from Rice Creek Field Station were female, with no males observed out of a sample of 170 individuals. Second, all four wild-caught adults, as well as the offspring of one of these adults, had identical genotypes at nearly all SNPs, consistent with clonal reproduction (Table 3). Third, the ratio of reference to alternate allele read counts in the sequencing data showed clear peaks at approximately 0.33 and 0.67 (Figure 1), consistent with triploid genotypes, which are known to be parthenogenic in European populations (Frankel 1979; Frankel et al. 1981; Fussey 1984). We are unable to rule out infection by sex-reversing endosymbionts such as *Wolbachia* in *T. pusillus*, since we did not perform PCR-based tests for its presence, but the observation of triploid genotypes is incompatible with typical diploid, meiotic sexual reproduction.

In contrast, *H. riparius* appears to be a typical sexual, diploid, outcrossing population. Although the sample size is limited due to the lower abundance of this species at our field site, the number of males and females sampled is not statistically different from a 50-50 sex ratio. Moreover, SNP data suggest that this *H. riparius* population is diploid (Figure 1) and outcrossing, with a much smaller proportion of SNPs showing identical genotypes across individuals (Table 3).

The observation that four randomly sampled *T. pusillus* individuals have identical genotypes at nearly all SNPs suggests that nearly all genetic diversity in this population is due to heterozygosity *within* individuals, rather than sequence differences *between* individuals. The lack of overall genetic differences among individuals suggests that all the *T. pusillus* specimens used for this experiment stem from the same genetic lineage. Although the number of sequenced individuals is low, if this observation holds up, it would be consistent with either a single, recent origin of parthenogenesis, or a population bottleneck during the introduction of this species into North America from its likely native Europe some time in the last several centuries (Jass & Klausmeier 2000). Studies on allozyme frequencies in European populations of *T. pusillus* have identified multiple distinct clones in European populations with independent origins of parthenogenesis (Christensen 1979; Friis Theisen et al. 1995), suggesting that the latter explanation for the low diversity among individuals in our population may be more likely.

*T. pusillus* displays much higher levels of heterozygosity than *H. riparius* (Table 4). These findings provide evidence that this triploid *T. pusillus* lineage may have arisen from a cross between two previously unknown divergent diploid lineages, similar to some other triploid parthenogens (Ament-Velásquez et al. 2016; Johnson & Howard 2007; Moritz 1983). An especially interesting parallel is the marbled crayfish, *Procambarus virginalis*, a triploid, parthenogenetic lineage first discovered in the aquarium trade in 1995, and now an invasive pest in the wild; genomic studies have found few differences across its distribution in the wild but much higher heterozygosity in parthenogenic, triploid individuals than in its diploid progenitor, leading to the hypothesis that it arose during a single mating between two distantly related diploids

(Gutekunst et al. 2018). Thus, *T. pusillus*, which was probably introduced into North America at least a few centuries ago like most common terrestrial isopods in the United States (Jass & Klausmeier 2000), may be an ideal model for studies of how invasive, parthenogenic species diversify and adapt as they spread and become established in new areas.

### Purifying selection in asexual lineages

Patterns of dN, dS, and dN/dS were not consistent with the expectation of relaxed purifying selection in asexual lineages (Figure 2). Although median dN and dS estimates were significantly higher in the parthenogenic *T. pusillus* than in *H. riparius*, these differences were small in magnitude and the distributions (among genes in each species) were almost entirely overlapping. Moreover, the median estimated dN/dS ratio was significantly lower in *T. pusillus*, though again the difference was small and the distributions of individual dN/dS estimates across genes overlapped substantially. Even if this difference is meaningful, it is in the opposite direction from theoretical predictions; relaxed purifying selection should result in higher dN/dS ratios in the parthenogenic *T. pusillus* lineage. Thus, between-species comparisons provide no evidence of inefficient purifying selection in this asexual lineage as predicted by theory and confirmed in some empirical studies (Hollister et al. 2015; Neiman et al. 2010; Johnson & Howard 2007; Henry et al. 2012; Bast et al. 2018; Maldonado et al. 2022). However, this may be due to the relatively young age of the asexual *T. pusillus* lineage as there would not have been ample time for nonsynonymous mutations to accumulate. This is especially important because, without recombination or gene conversion, homologous chromosomes will evolve independently, and novel mutations are not expected to become fixed within asexual species, but instead may persist as heterozygous SNPs, in what is known as the Meselson effect (Mark Welch & Meselson 2000; Ament-Velásquez et al. 2016). Thus, comparisons of assembled transcript sequences may not reveal the full scope of divergence between species, if heterozygous transcripts are “collapsed” into a single contig in a transcriptome assembly.

To account for this issue, we also examined pN/pS, the ratio of non-synonymous to nonsynonymous polymorphisms within species, in *T. pusillus* and *H. riparius*. Again, considering all SNPs in each species, we found no significant differences between asexual and sexual lineages (Table 4); if anything, pN/pS ratios are slightly lower in the asexual *T. pusillus*, though this difference is not significant, again counter to theoretical predictions. These results are also not necessarily surprising because, if the asexual *T. pusillus* lineage arose with a cross between two divergent diploid *T. pusillus* lineages, as suggested by its high heterozygosity, then the majority of the heterozygous SNPS in the asexual lineage would have originated earlier, as differences between the two divergent diploid lineages.

Finally, we examined pN/pS ratios considering only the subset of SNPs in *T. pusillus* that may have arisen recently, through new mutations after the transition to parthenogenesis. In this subset of SNPs, pN/pS ratios were significantly higher compared to *T. pusillus* SNPs where all individuals were identical, or compared to all SNPs in *H. riparius* (Table 4). This suggests that nonsynonymous mutations are indeed purged more quickly in the sexual *H. riparius*, supporting the hypothesis that inefficient purifying selection may impose an evolutionary cost on asexual lineages, at least over the long term.

We propose that the inconsistent results in these types of studies may be the result of variability in both the biology of the study species and the approaches used. For instance, evolutionarily older asexual lineages will have had more time for novel mutations to accumulate. Likewise, the mode of parthenogenesis is also expected to impact the rate at which new polymorphisms become homozygous in asexual lineages; for example, in apomictic species, heterozygous SNPs will be maintained across generations (except for presumably rare gene conversion events), whereas in automictic species, heterozygous mutant alleles can become homozygous more readily (Jaron et al. 2021). This difference will impact the derived allele’s “exposure” to selection, as well as whether it is detected in, say, a transcriptome-based molecular evolution study. For instance, if a transcriptome assembler randomly selects one allele as the “reference” allele during the assembly process, and only the “reference” contig sequences are used in dN/dS analyses, up to half of all novel mutations in an asexual lineage may be missed, reducing the statistical power of downstream analyses. And while some studies have considered both within- and between-species sequence variation (Johnson & Howard 2007; Bast et al. 2018), this is not universal (Brandt et al. 2017, 2019).

### Conclusions

This study set out to identify the reproductive mode of *T. pusillus* and *H. riparius* populations located in Oswego, NY as well as test the hypothesis that nonsynonymous mutations accumulate more rapidly in asexually reproducing organisms. We confirmed that this population of *T. pusillus* is asexual and triploid, and that the sampled population of *H. riparius* is sexual. Although patterns of molecular evolution were inconsistent with the hypothesis of inefficient purifying selection in asexual species when considering between-species divergence or all SNPs overall, we did find evidence of an elevated pN/pS ratio for putatively recent SNPs. These findings suggest that purifying selection on new mutations is less effective in asexual lineages, but that this effect may only become apparent across longer time scales. Our results indicate that future studies on the molecular evolution of asexual and sexual lineages should carefully consider both the biology of the particular study system (e.g., age of the asexual lineage, apomixis vs. automixis) as well as methodology for sequence analyses (e.g., accounting for SNPs instead of using only “reference” assembled sequences). We also propose that *T. pusillus*, including studies with wider geographic sampling, should provide a promising novel system for future studies on the evolutionary tradeoffs between sexual and asexual reproduction, given its distribution in North America and Europe and the occurrence of both sexual and asexual forms in the wild, particularly in the context of adaptation and diversification during introductions and range expansions.

## Methods

### Field Surveys and Identification of Species and Sex

All specimens were obtained from Rice Creek Field Station in Oswego, NY, during the months of June, July, and August 2021. Specimens were collected between 8:00 AM and 9:30 AM, and the same areas were sampled a total of four times over three months. To catch specimens, the bases of trees and fallen logs were surveyed and leaf litter was overturned. Homemade mouth aspirators were used to gather specimens from the ground and keep them in a secure container before they were transported to the lab.

To determine the species and sex of specimens, they were observed under a stereomicroscope. Each individual was anesthetized in a 0.01% clove oil solution, since their small size (adults are only a few mm long) and rapid movement make them difficult to handle and examine. Clove oil has been used to anesthetize aquatic isopods in other studies (Mojaddidi et al. 2018), and has previously been used to successfully anesthetize larger terrestrial isopods in our lab.

Specimens were identified to species following taxonomic keys (Shultz 2018; Van Name 1936). Although these two species appear superficially similar, the number of ocelli can be used to distinguish them: each *T. pusillus* eye has three ocelli, while *H. riparius* only has one (Shultz 2018; Van Name 1936). This was determined under the microscope by turning each specimen on its side to observe their eyes before moving on to sexing the specimens. To determine the sex of each isopod, the shape of the pleon was observed, with females having rounder scale rows and males having more pointed scale rows (Shultz 2018). Females were also identified through the obvious presence of brood pouches, as many were ovigerous during the study period.

After species identification and sexing, each specimen was placed in an individual deli cup with a substrate of moist soil and leaf litter along with thinly sliced carrots. Each specimen was kept separate from others to prevent the spread of any potential pathogens. This also allowed us to obtain full-sibling groups of offspring from females that were gravid at the time of capture.

### RNA Extractions

RNA extraction was performed on five *T. pusillus* specimens, including four randomly chosen wild-caught adult females and a pooled set of offspring from one of these adult females, and four *H. riparius* specimens, two males and two females. Prior to RNA extraction, the specimens were suspended in RNA Shield (Zymo Research) and stored at -20°C to preserve RNA. An NEB Monarch RNA Extraction Kit (New England BioLabs) was used according to the manufacturer’s instructions. RNA yield and quality were assessed using a NanoDrop and Qubit before being sent to a lab at University at Buffalo (UB) for sequencing.

### Data cleaning & trimming

Raw sequence reads were trimmed and filtered using the bbduk tool of the bbmap package (Bushnell, B) with the options ktrim=r, k=23, mink=11, hdist=1, tpe, tbo for adapter trimming.

### Transcriptome assembly

Following best practices (Gilbert 2013, 2019), we assembled transcriptomes using multiple assemblers and subsequently merged and filtered the assemblies to obtain a final assembly for each species. For each species, we separately assembled the transcriptome of one male and one female sample, each using Trinity v. 2.14.0 (Grabherr et al. 2011), TransLiG v. 1.3 (Liu et al. 2019), SOAPdenovo-trans v. 1.0.5 (Xie et al. 2014) with k=31, and RNASpades v. 3.15.4 (Bushmanova et al. 2019) using k=31. We then merged the 8 draft assemblies of each species (4 male assemblies plus 4 female assemblies) and filtered the merged assemblies using EvidentialGenes v. 3.18 (Gilbert 2013, 2019). After filtering, EvidentialGenes produces two sets of transcripts for each assembly: “main” transcripts, and a set of “alternate” transcripts, which are predicted to be alternative splicing variants. Next, we removed transcripts/contigs likely originating from contamination, such us bacteria and other symbionts. To accomplish this, we used BLAST+ v. 2.13.0 (Camacho et al. 2009) and diamond v. 2.0.15.153 (Buchfink et al. 2021) to look for hits in the NCBI nt and nr databases, respectively, using E-value threshold of 10^−25^ or better. Any transcript with a best hit originating from any taxon outside multicellular animals was discarded as a potential contaminant (transcripts with no hits were retained). Because our samples were multiplexed and sequenced together, we also ran CroCo v. 1.1 (Simion et al. 2018) to remove transcripts that may represent cross-contamination between species (e.g., due to index hopping).

The resulting transcriptome assemblies were assessed using BUSCO v. 5.3.2 (Manni et al. 2021) with the arthropoda odb10 gene set.

### SNP calling and dN/dS analyses

Trimmed sequence reads were aligned to the assembled transcriptomes using bwa-mem-2-v.2.3.2 (Vasimuddin et al. 2019). Sambamba v. 0.8.2 and samtools v. 1.15.1 were used to index and sort the resulting bam files, and filter out discordantly mapped sequence read pairs (Tarasov et al. 2015; Danecek et al. 2021). Finally, SNPs were called using freebayes-1.3.6 (Garrison & Marth 2012) with the specifications of a minimum mapping quality of 60, minimum coverage of 50, and ploidy being 2 for *H. riparius*, which is diploid, and 3 for *T. pusillus*, which our analyses showed is triploid. At the mapping stage, sequence reads were aligned to the full transcriptome assembly for each species (i.e., predicted main and alternate transcript sequences); however, SNPs were called only in the main transcripts, to avoid pseudo-replication caused by SNPs found in multiple transcript isoforms. In downstream analyses, we also retained only SNPs at which every sequenced individual had a sequencing depth of 20 or greater, a genotype quality score of 30 or greater, and where the SNP site overall had a quality score of 30 or greater, as estimated by freebayes. When computing SNP frequencies in each species, i.e., number of polymorphic sites / number of total sites sequenced, the total number of sites (in other words, the denominator) included only sites with a sequencing depth of at least 20 in each sequenced individual, because sites with low sequencing depth might appear monomorphic even if there is a true SNP at that location.

We predicted full-length protein sequences in each assembled transcriptome using Codan v. 1.1 (Nachtigall et al. 2021). We then used the predicted coding regions to estimate rates of nonsynonymous and synonymous polymorphism within each species in protein-coding regions (pN, pS, and pN/pS). To identify orthologs across all three species, we used OrthoFinder v. 2.5.4 (Emms & Kelly 2019). To estimate nonsynonymous and synonymous divergence across species (dN, dS, and dN/dS) we aligned predicted amino acid sequences of single-copy orthologs using MUSCLE v. 5.1 (Edgar 2004), back-converted amino acid alignments to nucleotide alignments using pal2nal v. 14.0 (Suyama et al. 2006) and then used the codeml module of PAML v. 4.9 (Yang 2007). We used the free ratios branch model (model=1) and performed pairwise comparisons (runmode=-2) between *T. pusillus* and *T. rathkei*, and between *H. riparius* and *T. rathkei*. Codon frequencies were used as free parameters (CodonFreq=3).

## Acknowledgments

We thank the editor and anonymous reviewers for helpful comments on earlier versions of this manuscript. We also appreciate computing resources provided by the Biomedical & Health Informatics program at SUNY Oswego and the NSF-supported Jetstream/Jetstream2 cloud computing platforms. This research was supported in part by National Science Foundation grant NSF-DEB 1453298 to CHC.

## Data Availability Statement

All sequence data generated for this project have been deposited at the NCBI Sequence Read Archive under BioProject Accession PRJNA916870 and SRA Accession Numbers SRR22938942-SRR22938950.

