## Supplementary Figure 1 for "Reduced effectiveness of purifying selection on new mutations in a parthenogenic terrestrial isopod (*Trichoniscus pusillus*)"

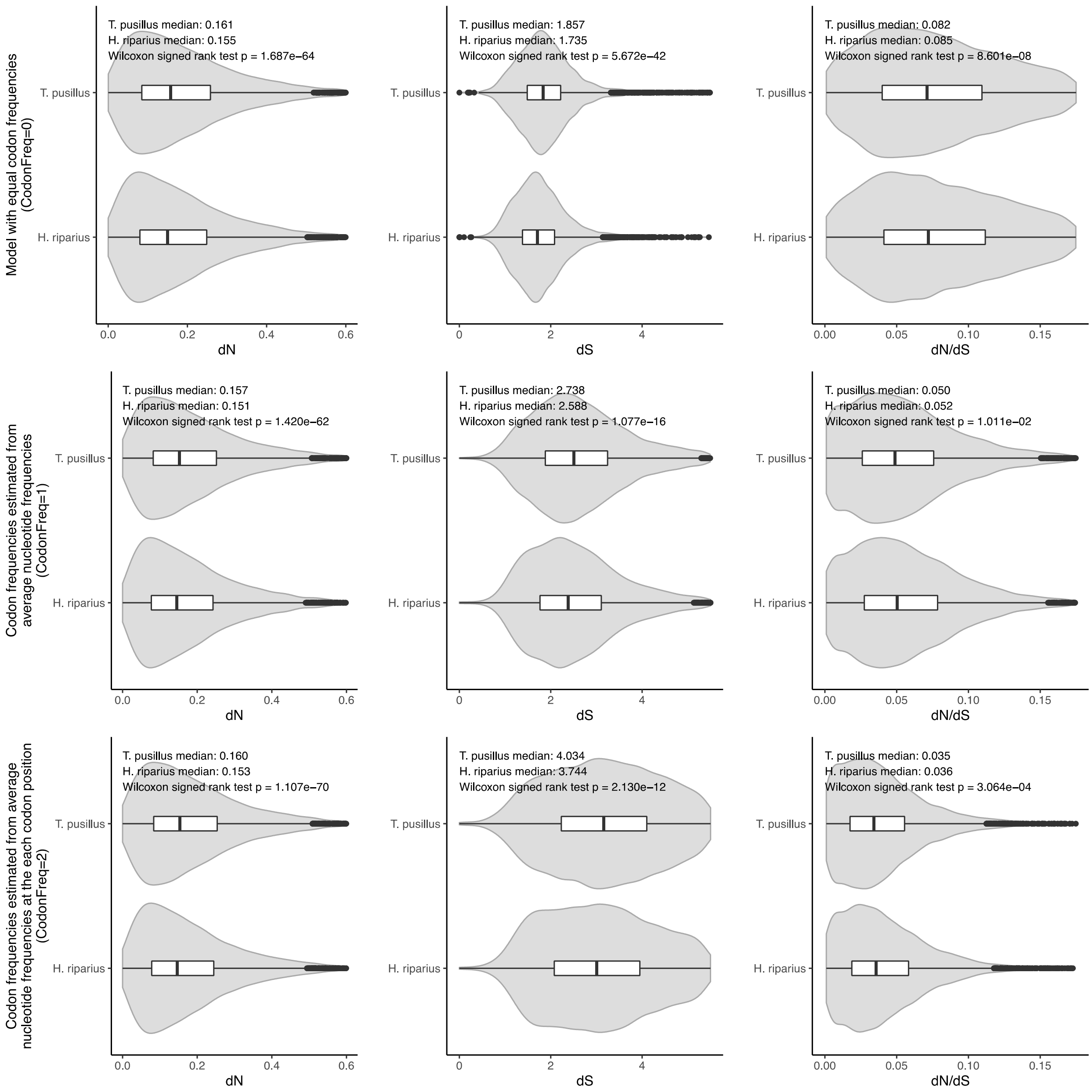


**Supplementary Figure 1.** Violin plots of *dN*, *dS*, and *dN/dS* ratios, resulting from pairwise comparisons between *Trichoniscus* vs. *Trachelipus*, and *Hyloniscus* vs. *Trachelipus*, using different codon models in the codeml package in PAML.
